# TP53-dependent CRISPR-Cas9 sex bias across cancer types affects MYC, PIK3CA, and SUV39H1 mediated by factors including SOX9, FOXO4, and PRC1

**DOI:** 10.1101/2022.08.03.502574

**Authors:** Mengbiao Guo, Yuanyan Xiong

## Abstract

CRISPR-Cas9 system has emerged as the dominant technology for gene editing and has great potential for large-scale clinical applications. One major concern is its off-target issue and other potential side effects after the introduction of exogenous CRISPR-Cas9 into cells. Several previous studies investigated CRISPR-Cas9 interactions with p53 mainly in non-transformed cells, such as RPE1 (retinal pigmented epithelium cells) and H9 (embryonic stem cells [ESC]). Recently, it has been reported that Cas9 alone can activate the p53 pathway and select for p53-inactivating mutations after studying hundreds of cancer cell lines. We reanalyzed the reported data of Cas9-associated p53-inactivating mutations and observed large significant sex difference when comparing Cas9 activities in p53-wildtype and p53-mutant cell lines. To expand the impact of this finding, we further examined all protein-coding genes screening by the CRISPR-Cas9 system in a large-scale dataset from the DepMap project. We highlight the p53 status-dependent sex bias of CRISPR-Cas9 effect across cancer cell types (genes including *MYC, PIK3CA, KAT2B, KDM4E, SUV39H1, FANCB, TLR7, and APC2*) and potential mechanisms (mediated by transcriptional factors including SOX9, FOXO4, LEF1, and RYBP) underlying this phenomenon, which suggest that the p53-dependent sex bias effect may need to be considered in future clinical applications, especially in cancer, when using this genome editing system.

## Main

CRISPR-Cas9, an emerging powerful genome-editing system originated form bacteria, is known to interact with p53[1-5]. This system can inactivate genes in a precise manner, relying on DNA breaks and repair, which are mostly guarded by p53. Enache *et al*.[6] have recently demonstrated that the protein Cas9 alone can activate the p53 pathway and select for p53-inactivated mutants. Briefly, they investigated the effects of Cas9 introduction on cells at the mRNA and protein levels, especially for the p53 pathway, and they further studied the differential Cas9 activities in cell lines with a wildtype (WT) or mutant (MUT) p53. They found that Cas9 introduction will activate the p53 pathway and select p53-inactivating mutations in examined cell lines. Interestingly, we found that the largest effects of many results shown by Enache *et al*. were from cell lines from female donors, including the largest fold-change of p53 activation (BT159: female), more DNA damage foci (MCF7: female), and the largest *TP53*-inactivating subclonal mutations expanding or shrinking (293T, HCC1419, and OVK18: female, **Fig. 1A**). In contrast, the difference of Cas9 activity between p53-WT (wildtype) and p53-MUT (mutant) cells was larger in male (two-sided t-test, *P*=3.1E-5, difference 14.85%) than female lines (two-sided Wilcoxon-test, *P*=0.021, difference 7.46%). The difference was much larger (*P*-value <2e-16, see **Methods**) in male compared with female cell lines (**Fig. 1B-C**). These observations clearly demonstrated significant sex bias of Cas9 functions in mammalian cells.

**Fig. 1.**
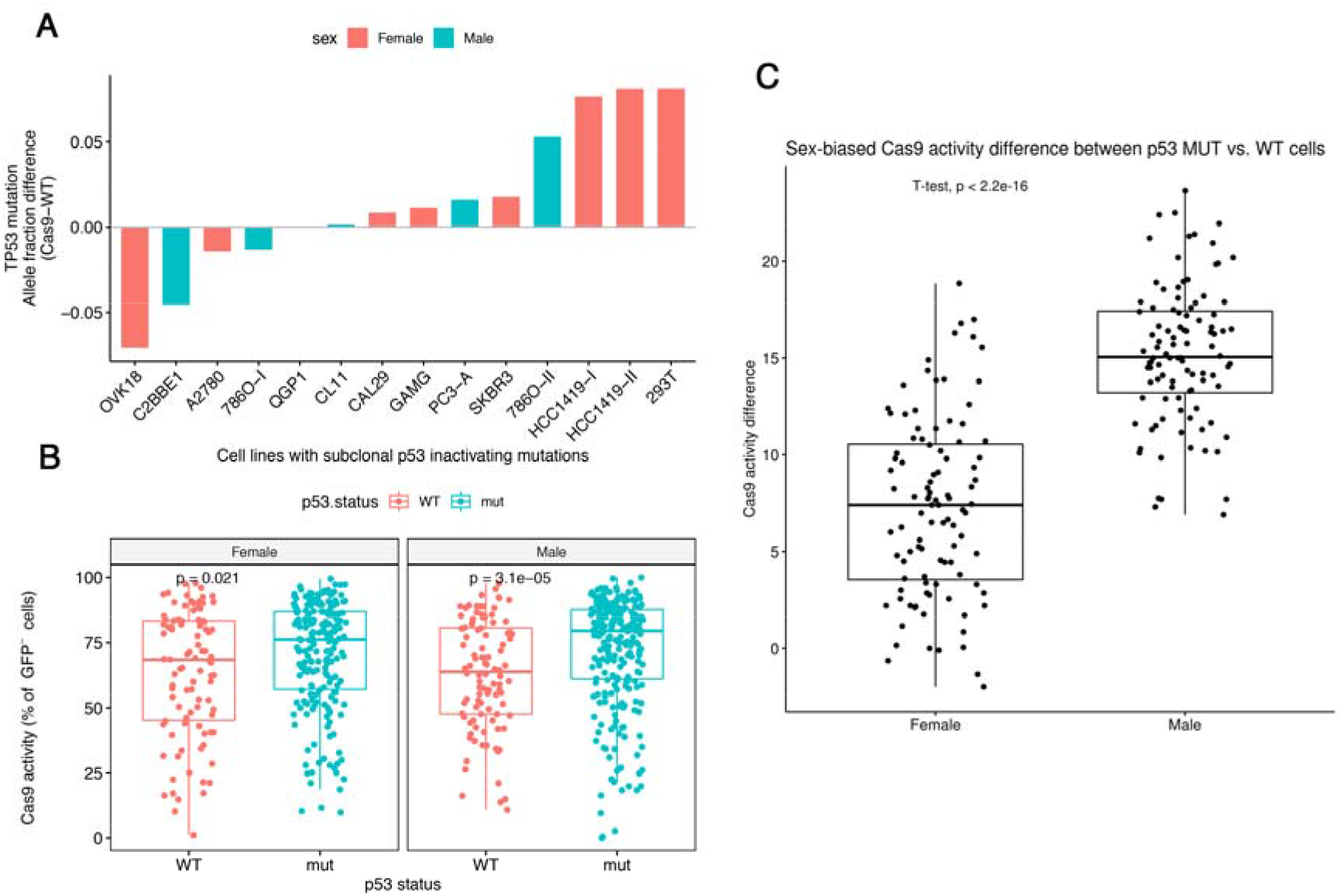
Sex-biased activities of Cas9 across cell lines are dependent on p53 status. (**A**) *TP53* allele fraction differences (y-axis) after Cas9 introduction in various cell lines (x-axis), colored by sexes. (**B**) Comparison of Cas9 activities (y-axis) between p53-WT and p53-MUT (x-axis) in male cell lines (right) and in female cell lines (left). Bar, median; box, 25th and 75th percentile; whiskers, 1.5 times the interquartile range of the lower and upper quartiles; points, individual cell lines. Two-sided Wilcoxon-test. (**C**) Box plots showing sex-biased (x-axis) Cas9 activity difference (y-axis) between p53-WT and p53-MUT cell lines shown in (**B**).

These results possibly reflect the sex-biased effects of Cas9 due to the differential functions and regulations of p53 between sexes. p53 is differentially regulated in both healthy and cancer samples between females and males[7]. Our recent work also shows that p53 regulates the active X chromosome dosage compensation, but with different effect sizes between sexes [8]. Importantly, some vital genomic regulators interacting with p53 are encoded by genes on chromosome X, which contributed to the sex bias of p53 activity, especially in cancer [7].

Since Cas9 is a key component of the CRISPR-Cas9 system, sex-biased activities of Cas9 itself can probably be extended to CRISPR-Cas9. Due to the potentially large-scale clinical usage of CRISPR-Cas9 in the future in patients with various diseases, sex-specific effects should be studied carefully and considered beforehand. However, we have found no report discussing sex-biased CRISPR-Cas9 knockout (KO) effects to date. We thus investigated the potential p53 status-dependent sex bias in the CRISPR screening data from the DepMap[9] project (22Q4, https://depmap.org/portal/). Our rational is that if the CRISPR-Cas9 KO effect of a gene shows significant sex bias and is p53 status-dependent, two scenarios may occur: (i) this sex difference can only be detected within either p53-WT or p53-MUT cell lines; (ii) or the sex difference can be detected in both p53-WT and p53-MUT cell lines but their magnitude differ significantly. This study only aimed to propose the concept of p53 status-dependent sex bias in CRISPR-Cas9 editing and provide basic evidence, therefore, the second scenario was not considered here because it was more complex, although we believe it may provide more significant genes and thus more insight.

First, we obtained a list of 65 genes showing significant p53-dependent sex-bias of KO effects (defined by the depletion of guide RNAs [gRNA] after CRISPR-Cas9 introduction into cells) by comparing CRISPR-Cas9 editing effect scores (calculated by the DepMap project) on genes between cell lines from male and female donors for each tissue (**Table 1, Table S1**). Several interesting genes were shown as examples, including DNASE1 (Deoxyribonuclease 1, female-biased), POLB (DNA Polymerase Beta, male-biased), DFFA (DNA Fragmentation Factor Subunit Alpha, male-biased), FANCB (Fanconi Anemia Group B Protein involved in DNA repair, female-biased), APC2 (Adenomatosis Polyposis Coli 2, female-biased), KAT2B (Histone Acetyltransferase PCAF, female-biased), and KDM4E (Lysine (K)-Specific Demethylase 4E, male-biased) (**Fig. 2A-G**) These genes included eight identified only based on genes (gene set 1) whose products with p53 protein-protein interactions (PPI, from the STRING database https://string-db.org/, see **Methods** for details), instead of all protein-coding genes (gene set 2), and another four identified based on both gene sets. Interestingly, only eight genes were located on the X chromosome, and only 13 out of 65 genes (**Table S1**), including H2AC16 (HIST1H2AL, H2A Clustered Histone 16, female-biased), APC2, DFFA, DNASE1, PRKRA (Protein Activator Of Interferon Induced Protein Kinase EIF2AK2/PKR, male-biased), KAT2B, FEN1 (Flap Structure-Specific Endonuclease 1, male-biased), were significant in p53-MUT samples. Moreover, we found sex-biased KO effects in both p53-WT and p53-MUT cells, but not for *TP53* (however, note that its negative regulator *MDM2* had a suggestive *P*-value of 0.0016 (two-sided t-test; male-biased) only in brain cells, which may help explain the p53 activation results in brain tumor BT159[6]. Of note, the FANCB of the Fanconi Anemia (FA) pathway has been identified to play a key role in CRISPR-Cas9 editing [10].

**Table 1.**
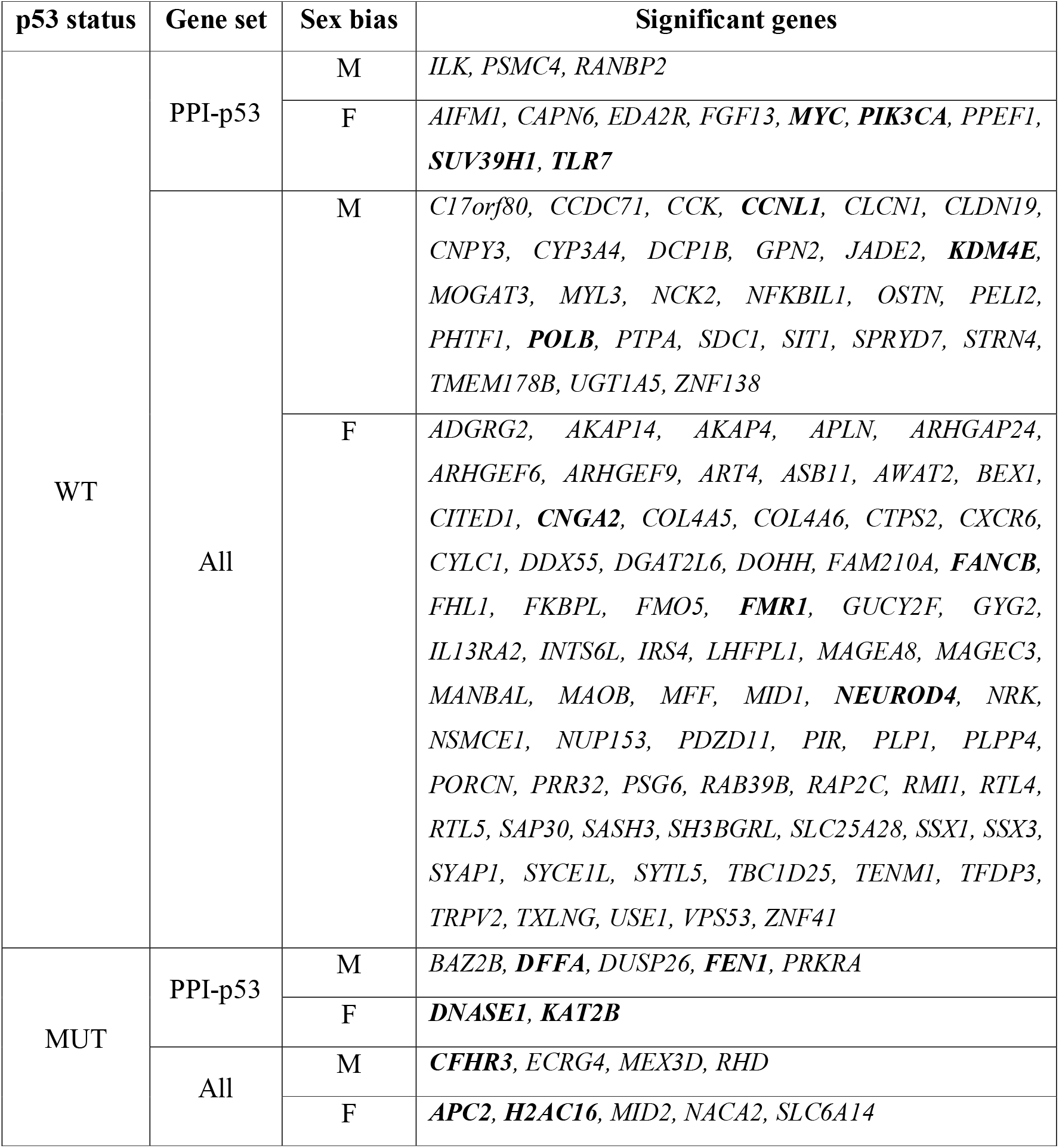
PCSB genes that show significantly sex-biased effects in CRISPR-Cas9 screening experiments. M: Male, F: Female.

**Fig. 2.**
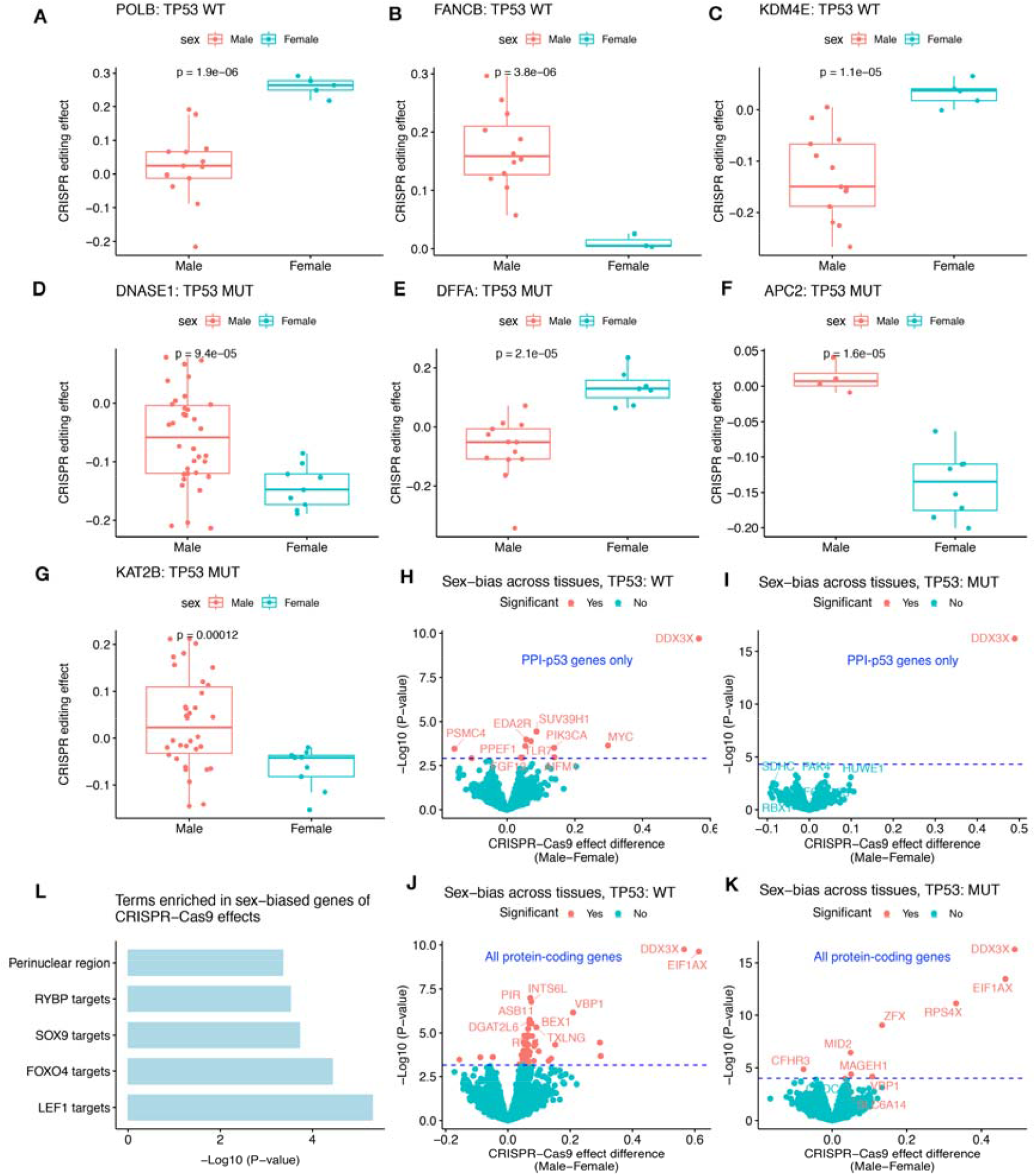
Sex-biased gene editing effects by CRISPR-Cas9 in cancer cells are dependent on p53 status. (**A-G**) Boxplots showing PCSB genes with sex-biased KO effects in CRISPR-Cas9 screening in a specific cancer cell type with (TP53-MUT) or without (TP53-WT) p53-inactivating mutation. Dots represent samples and were colored by sex. Two-sided t-test. (**H-K**) Volcano plots showing significantly sex-biased CRISPR-Cas9 KO effects (y-axis: *P*-values, x-axis: gene editing effect differences) across cancer types with TP53-MUT (**H, J**) or TP53-WT (**I, K**), based on genes encoding proteins interacting with p53 only or all protein-coding genes. The ten most promising genes to show sex bias, ranked by nominal *P*-values, were labeled. Dashed blue lines represent significance cutoff after multiple testing correction by false discovery rate (FDR). (**L**) Barplots showing the significance of enrichment in targets of four transcription factors (TFs, including RYBP, SOX9, FOXO4, and LEF1) or in perinuclear regions for p53-dependent sex-biased genes of CRISPR-Cas9 editing.

Moreover, we conducted a cross-tissue investigation of sex-biased gene editing effects after controlling for tissue types, which resulted in another 64 significant genes (CNGA2 [Cyclic Nucleotide Gated Channel Subunit Alpha 2] was also identified in single-tissue analysis above, female-biased), such as MYC (Myc Proto-Oncogene Protein, female-biased), SUV39H1 (SUV39H1 Histone Lysine Methyltransferase, also known as KMT1A, female-biased), FMR1 (Fragile X Messenger Ribonucleoprotein 1, female-biased), PIK3CA (PtdIns-3-Kinase Catalytic Subunit Alpha, female-biased), and TLR7 (Toll Like Receptor 7, female-biased) (**Fig. 2H-K, Table 1, Table S1**). Notably, only 3 out of 64 genes, MID2 (Midline 2, RING-Type E3 Ubiquitin Transferase), CFHR3 (Complement Factor H Related 3), and SLC6A14 (Solute Carrier Family 6 Member 14), were identified as significant in p53-MUT samples (**Fig. 2K**, note that DDX3X, EIF1AX, RPS4X, ZFX, MAGEH1, and VBP1 were excluded because they were identified as significant in both p53-WT and p53-MUT samples), consistent with the single-tissue analysis results above. In contrast to single-tissue analysis, 52 out of 64 genes were located on chromosome X (**Table S1**).

Interestingly, MYC is located on the X chromosome in Drosophila and its sex-specific dosage regulates body size differentially between females and males [11]. Also in Drosophila, the sex-determining gene daughterless has sequence similarities to MYC [12]. p53 can regulate MYC in various ways and vice versa [13-20]. Moreover, MYC function was controlled by the androgen receptor (AR)/p53 axis [20], and by DDX3Y male MYC-driven lymphomas with DDX3X-inactivation [21]. Additionally, MYC is a direct target of APC (WNT Signaling Pathway Regulator), the paralog of APC2 [22].

To further reveal the potential mechanism of p53 regulating Cas9 activities in human cells, we performed functional enrichment analysis for all these 128 significant genes with p53-dependent CRISPR-Cas9 sex bias (termed PCSB genes). We observed that they were enriched in perinuclear regions (p=5.0e-6, reported by the webserver of ToppGene [23], https://toppgene.cchmc.org/enrichment.jsp) and in target genes of four transcription factors (TF): LEF1 (p=3.7e-5, reported by the webserver of ToppGene; Lymphoid Enhancer Binding Factor 1), FOXO4 (p=1.9e-4; Forkhead Box O4), SOX9 (p=3.0e-4; SRY-Box Transcription Factor 9), and RYBP (p=4.4e-4; RING1 And YY1 Binding Protein) (**Fig. 2L**). We believe these TFs are potential drivers of the sex bias studied in this work.

Three of the four TFs were directly linked to sex or sex chromosomes. First, the gene encoding FOXO4 is located on chromosome X. FOXO4 has PPI interaction with p53 [24, 25] and also MDM2 (Mouse Double Minute 2, Human Homolog Of; P53-Binding Protein), the prime E3 ligase for p53 [26, 27]. Moreover, PIK3CA is the catalytic subunit of PI3K and FOXO4 is a downstream effector of the PI3K/AKT pathway [28]. Second, RYBP is a component of the Polycomb group (PcG) protein complex 1 (PRC1) that mediates the histone modification of H2AK119ub and is required for X chromosome inactivation (XCI) in females [29]. RYBP can stabilize p53 by modulating MDM2 [30]. Third, SOX9 on chromosome 17 is a direct target of SRY (Sex-Determining Region Y) and, together with its distal enhancer, is critical in mammalian sex determination [31]. SOX9 can regulate MYC expression by interacting with FOXC1 [32]. The remaining TF, LEF1, can be induced by MYC to activate the WNT pathway and sustain cell proliferation [33]. LEF1 can also activate MYC or CCND1 to enhance pancreatic tumor cell proliferation [34, 35].

Of note, p53 can negatively regulate SUV39H1 to remodel H3K9me3-marked constitutive heterochromatin by recruiting HP1 [36, 37], while SUV39H1 is functionally linked with PRC1 [38]. MYC is also an important interactor with another PRC1 subunit YAF2 (YY1 Associated Factor 2) [39]. Consistently, a recent study reported higher activity of CRISPR-Cas9 in active chromatin than in heterochromatin regions [40], therefore, it is possible that p53-dependent sex-biased CRISPR-Cas9 activity was mediated by chromatin remodelers, such as PRC1, SUV39H1, KAT2B, and KDM4E.

Finally, a recent report reported both CRISPR-specific differentially essential positive (CDE+) and CDE-genes dependent on TP53 or KRAS [41]. We found that PCSB genes were in both TP53-dependent CDE+ genes (CCNL1, FAM210A, MOGAT3, and FKBPL) and CDE-genes (NEUROD4), and in KRAS-dependent CDE-genes (PELI2, NACA2, and RMI1) reported by Sinha et al [41]. More importantly, we have detected sex bias in genes of potential CRISPR-selected cancer drivers (CCD) similar to TP53 from the same study [41]. These genes include PIK3CA (FDR=3.2e-29 as reported by Sinha et al.) and the paralog of APC2 (APC, FDR=0.1) from our PCSB genes. Interestingly, SOX9 (FDR=2.5e-9 as reported by Sinha et al., one of the four TFs with targets enriched in our PCSB genes) was also among their CCD genes.

We summarized our findings and proposed potential mechanisms that p53 can affect CRISPR-Cas9 effects in mammalian cells (**Fig. 3**). Collectively, our work demonstrate the potential sex-biased gene editing effects both at the genome-wide level based on Cas9 itself and for specific genes based on CRISPR-Cas9 screening results. We suggest that the sex bias effect may need to be considered in future clinical applications (e.g. targeting MYC, PIK3CA, APC2, FANCB, KAT2B, KDM4E, TLR7, and SUV39H1 from our PCSB genes), especially in cancer, when using CRISPR-Cas9 and potentially other Cas protein systems.

**Fig. 3.**
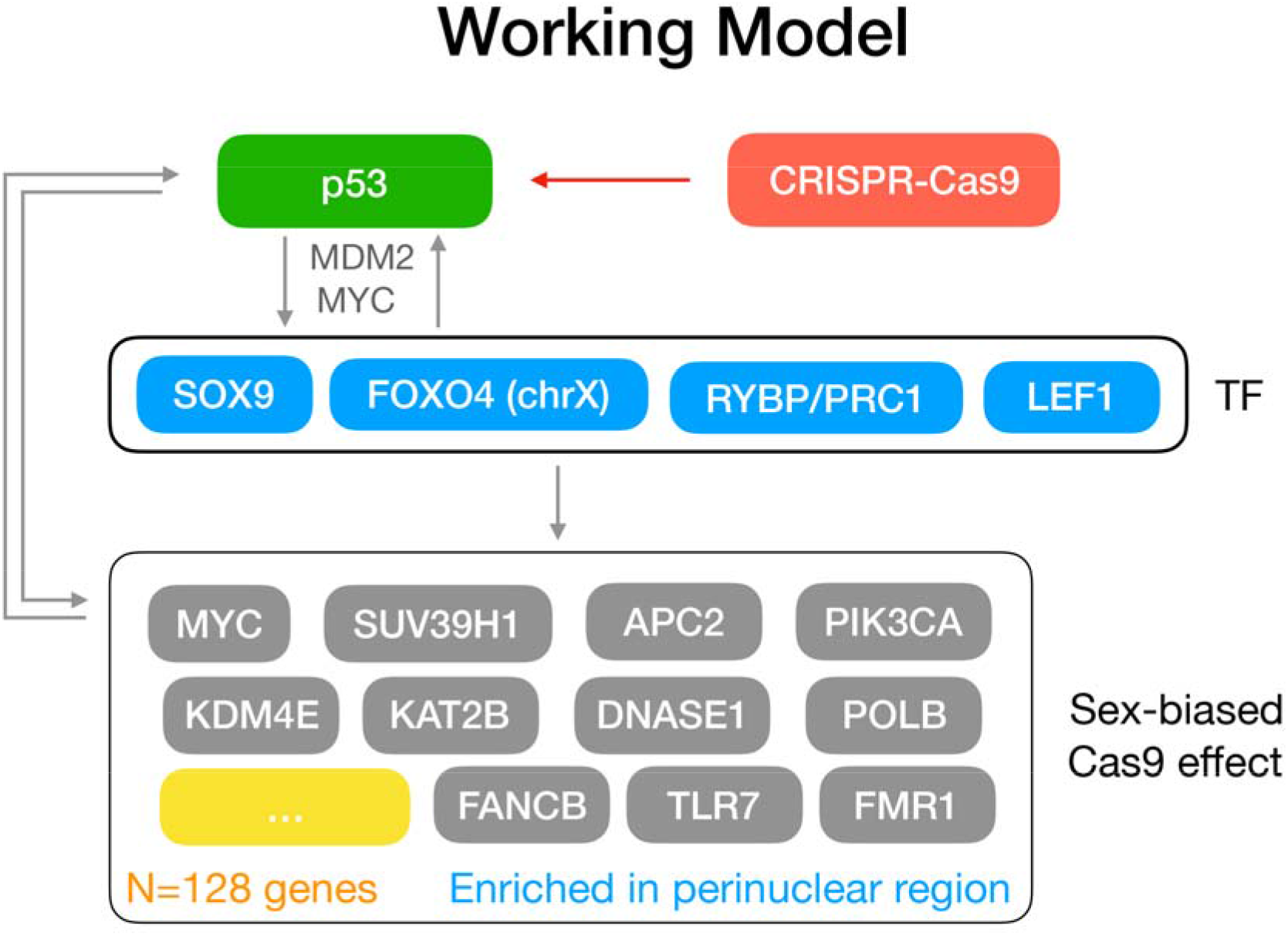
The proposed working model showing potential mechanisms p53 status can affect CRISPR-Cas9 editing effects differently between sexes in mammalian cells. p53 may affect CRISPR-Cas9 in a sex-dependent way either directly by regulating some of the sex-biased genes or indirectly by regulating four TFs, SOX9, FOXO4, RYBP, or LEF1, which may in turn be mediated by MDM2 or MYC.

## Methods

We obtained processed data of gene editing KO effects for CRISPR screening, gene mutations from Cancer Cell Line Encyclopedia (CCLE), and all cell line information from the DepMap project (22Q4, https://depmap.org/portal/). CRISPR KO effects were defined by the depletion of guide RNAs (gRNA) after CRISPR-Cas9 introduction into cell lines followed by sequencing and were calculated by comparing gRNA numbers in the initial cells at the beginning with those in the cells cultured for a period of time. Cell lines (excluded if with ‘unknown’ sex information) were separated into two groups by p53 status, p53-WT and p53-MUT (with p53 non-silent mutations as defined in the Enache *et al* study). For both p53 groups, cell lineages with >=10 samples (at least 3 samples for both males and females) were retained for downstream analysis. First, we tested the sex-biased KO effects in each cell lineage for a list of 2,007 genes (termed ‘gene set 1’) encoding proteins interacting with p53, obtained from the PPI database STRING (https://string-db.org/) by requiring interaction confidence scores larger than 400. Then, we extended the test to all genes (n=17,454, termed ‘gene set 2’) screened by DepMap. Differential KO effects between sexes for each tissue were compared by two-sided t-test. Cross-tissue sex-biased KO effects were obtained by applying limma [42] to KO effects of all cell lines (excluding those from sex-specific tissues: breast, cervix, ovary, prostate, uterus) at the same time, with sex and tissue type as covariables.

Of note, seven genes (EDA2R, MYC, PIK3CA, PPEF1, PSMC4, SUV39H1, and TLR7) were identified as significant based on both ‘gene set 1’ and ‘gene set 2’, and only annotated as genes whose protein products with p53 PPI. Moreover, five genes (DDX3X, EIF1AX, RPS4X, ZFX, MAGEH1, and VBP1) showed significant CRISPR-Cas9 sex bias in both p53-WT and p53-MUT samples and were excluded. Among them, DDX3X, EIF1AX, and ZFX are functionally linked with their paralogs encoded by the Y chromosome in males (DDX3Y, EIF1AY, and ZFY, respectively) [43, 44] and escape X inactivation in females [45]. The potential loss of chromosome Y in some male samples or reactivation of chromosome X in some female samples may explain these results.

Throughout this study, *P*-values were corrected for multiple testing by the Benjamini & Hochberg (or false discovery rate, FDR) correction [46] by using the R function p.adjust. Adjusted *P*-values smaller than 0.2 were considered significant and raw *P*-values less than 2e-3 were considered suggestive. All analysis and visualizations were performed using R language (http://www.r-project.org/).

Functional enrichment analysis of significant genes with sex bias of CRISPR-Cas9 editing (PCSB genes) was performed by using the web server of ToppGene (https://toppgene.cchmc.org/enrichment.jsp) [23].

Number of cell lines for each tissue used in the CRISPR-Cas9 analysis were as following: bile_duct (n=20), blood (n=49), bone (n=25), central_nervous_system (n=66), colorectal (n=46), esophagus (n=30), eye (n=7), gastric (n=27), kidney (n=20), liver (n=22), lung (n=125), lymphocyte (n=34), pancreas (n=41), peripheral_nervous_system (n=24), plasma_cell (n=18), skin (n=52), soft_tissue (n=27), thyroid (n=13), upper_aerodigestive (n=51), urinary_tract (n=28). The subgroup numbers regarding to p53 status (WT or MUT) and sex (male or female) were shown in **Table S2**.

Bootstrapping was used to assess significance of the sex-bias of Cas9 activity differences between MUT and WT p53 cell lines. A total of 100 random samples were obtained from Cas9 activities of cell lines, and Cas9 differences for each sex between MUT and WT p53 were calculated for each random sample.

## Data and code availability

All data sets are available within the article or its Supplementary Information. DepMap data were downloaded from https://depmap.org/. Protein-protein interactions with *TP53* were downloaded from STRING (http://string-db.org/).

## Acknowledgement

We thank the DepMap and CCLE projects for making their datasets available for analysis. The research has been supported by Integrated Project of Major Research Plan of National Natural Science Foundation of China (NSFC) (Grant 92249303) and Guangdong Basic and Applied Basic Research Foundation (2021A1515110972).

## Author contributions

MBG conceived the idea, performed the analysis, and drafted the manuscript. YYX provided writing input. Both authors reviewed the manuscript.

## Supplementary Table Legends

**Table S1**. Genes with p53-dependent sex-biased effects in CRISPR-Cas9 screening.

**Table S2**. Number of cell lines regarding p53 status and sex.

## References

1. Ihry, R.J., et al., p53 inhibits CRISPR-Cas9 engineering in human pluripotent stem cells. Nat Med, 2018. 24(7): p. 939–946.

2. Haapaniemi, E., et al., CRISPR-Cas9 genome editing induces a p53-mediated DNA damage response. Nat Med, 2018. 24(7): p. 927–930.

3. Wu, Y., et al., Highly efficient therapeutic gene editing of human hematopoietic stem cells. Nat Med, 2019. 25(5): p. 776–783.

4. Schiroli, G., et al., Precise Gene Editing Preserves Hematopoietic Stem Cell Function following Transient p53-Mediated DNA Damage Response. Cell Stem Cell, 2019. 24(4): p. 551–565 e8.

5. Brown, K.R., et al., CRISPR screens are feasible in TP53 wild-type cells. Mol Syst Biol, 2019. 15(8): p. e8679.

6. Enache, O.M., et al., Cas9 activates the p53 pathway and selects for p53-inactivating mutations. Nat Genet, 2020.

7. Haupt, S., et al., Identification of cancer sex-disparity in the functional integrity of p53 and its X chromosome network. Nat Commun, 2019. 10(1): p. 5385.

8. Guo, M., et al., Distinct dosage compensations of ploidy-sensitive and -insensitive X chromosome genes during development and in diseases. iScience, 2023. 26(2): p. 105997.

9. Meyers, R.M., et al., Computational correction of copy number effect improves specificity of CRISPR-Cas9 essentiality screens in cancer cells. Nat Genet, 2017. 49(12): p. 1779–1784.

10. Richardson, C.D., et al., CRISPR-Cas9 genome editing in human cells occurs via the Fanconi anemia pathway. Nat Genet, 2018. 50(8): p. 1132–1139.

11. Mathews, K.W., M. Cavegn, and M. Zwicky, Sexual Dimorphism of Body Size Is Controlled by Dosage of the X-Chromosomal Gene Myc and by the Sex-Determining Gene tra in Drosophila. Genetics, 2017. 205(3): p. 1215–1228.

12. Caudy, M., et al., daughterless, a Drosophila gene essential for both neurogenesis and sex determination, has sequence similarities to myc and the achaete-scute complex. Cell, 1988. 55(6): p. 1061–7.

13. Feng, Y.C., et al., c-Myc inactivation of p53 through the pan-cancer lncRNA MILIP drives cancer pathogenesis. Nat Commun, 2020. 11(1): p. 4980.

14. Petr, M., et al., Wild-type p53 binds to MYC promoter G-quadruplex. Biosci Rep, 2016. 36(5).

15. Olivero, C.E., et al., p53 Activates the Long Noncoding RNA Pvt1b to Inhibit Myc and Suppress Tumorigenesis. Mol Cell, 2020. 77(4): p. 761–774 e8.

16. Sachdeva, M., et al., p53 represses c-Myc through induction of the tumor suppressor miR-145. Proc Natl Acad Sci U S A, 2009. 106(9): p. 3207–12.

17. Morcelle, C., et al., Oncogenic MYC Induces the Impaired Ribosome Biogenesis Checkpoint and Stabilizes p53 Independent of Increased Ribosome Content. Cancer Res, 2019. 79(17): p. 4348–4359.

18. Liao, P., et al., Mutant p53 Gains Its Function via c-Myc Activation upon CDK4 Phosphorylation at Serine 249 and Consequent PIN1 Binding. Mol Cell, 2017. 68(6): p. 1134–1146 e6.

19. Lindstrom, M.S. and K.G. Wiman, Myc and E2F1 induce p53 through p14ARF-independent mechanisms in human fibroblasts. Oncogene, 2003. 22(32): p. 4993–5005.

20. Cottle, D.L., et al., c-MYC-induced sebaceous gland differentiation is controlled by an androgen receptor/p53 axis. Cell Rep, 2013. 3(2): p. 427–41.

21. Gong, C., et al., Sequential inverse dysregulation of the RNA helicases DDX3X and DDX3Y facilitates MYC-driven lymphomagenesis. Mol Cell, 2021. 81(19): p. 4059–4075 e11.

22. He, T.C., et al., Identification of c-MYC as a target of the APC pathway. Science, 1998. 281(5382): p. 1509–12.

23. Chen, J., et al., ToppGene Suite for gene list enrichment analysis and candidate gene prioritization. Nucleic Acids Res, 2009. 37(Web Server issue): p. W305–11.

24. Mandal, R., et al., FOXO4 interacts with p53 TAD and CRD and inhibits its binding to DNA. Protein Sci, 2022. 31(5): p. e4287.

25. Baar, M.P., et al., Targeted Apoptosis of Senescent Cells Restores Tissue Homeostasis in Response to Chemotoxicity and Aging. Cell, 2017. 169(1): p. 132–147 e16.

26. Brenkman, A.B., et al., Mdm2 induces mono-ubiquitination of FOXO4. PLoS One, 2008. 3(7): p. e2819.

27. Oteiza, A. and N. Mechti, The human T-cell leukemia virus type 1 oncoprotein tax controls forkhead box O4 activity through degradation by the proteasome. J Virol, 2011. 85(13): p. 6480–91.

28. Brunet, A., et al., Akt promotes cell survival by phosphorylating and inhibiting a Forkhead transcription factor. Cell, 1999. 96(6): p. 857–68.

29. Hernandez-Munoz, I., et al., Stable X chromosome inactivation involves the PRC1 Polycomb complex and requires histone MACROH2A1 and the CULLIN3/SPOP ubiquitin E3 ligase. Proc Natl Acad Sci U S A, 2005. 102(21): p. 7635–40.

30. Chen, D., et al., RYBP stabilizes p53 by modulating MDM2. EMBO Rep, 2009. 10(2): p. 166–72.

31. Gonen, N., et al., Sex reversal following deletion of a single distal enhancer of Sox9. Science, 2018. 360(6396): p. 1469–1473.

32. Tang, L., et al., SOX9 interacts with FOXC1 to activate MYC and regulate CDK7 inhibitor sensitivity in triple-negative breast cancer. Oncogenesis, 2020. 9(5): p. 47.

33. Hao, Y.H., et al., Induction of LEF1 by MYC activates the WNT pathway and maintains cell proliferation. Cell Commun Signal, 2019. 17(1): p. 129.

34. Jesse, S., et al., Lef-1 isoforms regulate different target genes and reduce cellular adhesion. Int J Cancer, 2010. 126(5): p. 1109–20.

35. Lim, S.K. and F.M. Hoffmann, Smad4 cooperates with lymphoid enhancer-binding factor 1/T cell-specific factor to increase c-myc expression in the absence of TGF-beta signaling. Proc Natl Acad Sci U S A, 2006. 103(49): p. 18580–5.

36. Park, J.W. and Y.S. Bae, Dephosphorylation of p53 Ser 392 Enhances Trimethylation of Histone H3 Lys 9 via SUV39h1 Stabilization in CK2 Downregulation-Mediated Senescence. Mol Cells, 2019. 42(11): p. 773–782.

37. Zheng, H., et al., p53 promotes repair of heterochromatin DNA by regulating JMJD2b and SUV39H1 expression. Oncogene, 2014. 33(6): p. 734–44.

38. Puschendorf, M., et al., PRC1 and Suv39h specify parental asymmetry at constitutive heterochromatin in early mouse embryos. Nat Genet, 2008. 40(4): p. 411–20.

39. Madge, B., et al., Yaf2 inhibits Myc biological function. Cancer Lett, 2003. 193(2): p. 171–6.

40. Alvarez, M.M., J. Biayna, and F. Supek, TP53-dependent toxicity of CRISPR/Cas9 cuts is differential across genomic loci and can confound genetic screening. Nat Commun, 2022. 13(1): p. 4520.

41. Sinha, S., et al., A systematic genome-wide mapping of oncogenic mutation selection during CRISPR-Cas9 genome editing. Nat Commun, 2021. 12(1): p. 6512.

42. Ritchie, M.E., et al., limma powers differential expression analyses for RNA-sequencing and microarray studies. Nucleic Acids Res, 2015. 43(7): p. e47.

43. Koferle, A., et al., Interrogation of cancer gene dependencies reveals paralog interactions of autosome and sex chromosome-encoded genes. Cell Rep, 2022. 39(2): p. 110636.

44. Venkataramanan, S., et al., DDX3X and DDX3Y are redundant in protein synthesis. RNA, 2021. 27(12): p. 1577–1588.

45. Navarro-Cobos, M.J., B.P. Balaton, and C.J. Brown, Genes that escape from X-chromosome inactivation: Potential contributors to Klinefelter syndrome. Am J Med Genet C Semin Med Genet, 2020. 184(2): p. 226–238.

46. Benjamini, Y. and Y. Hochberg, Controlling the False Discovery Rate: A Practical and Powerful Approach to Multiple Testing. Journal of the Royal Statistical Society: Series B (Methodological), 1995. 57(1): p. 289–300.

